# AITL: Adversarial Inductive Transfer Learning with input and output space adaptation for pharmacogenomics

**DOI:** 10.1101/2020.01.24.918953

**Authors:** Hossein Sharifi-Noghabi, Shuman Peng, Olga Zolotareva, Colin C. Collins, Martin Ester

## Abstract

**Motivation:** The goal of pharmacogenomics is to predict drug response in patients using their single- or multi-omics data. A major challenge is that clinical data (i.e. patients) with drug response outcome is very limited, creating a need for transfer learning to bridge the gap between large pre-clinical pharmacogenomics datasets (e.g. cancer cell lines), as a source domain, and clinical datasets as a target domain. Two major discrepancies exist between pre-clinical and clinical datasets: 1) in the input space, the gene expression data due to difference in the basic biology, and 2) in the output space, the different measures of the drug response. Therefore, training a computational model on cell lines and testing it on patients violates the i.i.d assumption that train and test data are from the same distribution.

**Results:** We propose Adversarial Inductive Transfer Learning (AITL), a deep neural network method for addressing discrepancies in input and output space between the pre-clinical and clinical datasets. AITL takes gene expression of patients and cell lines as the input, employs adversarial domain adaptation and multi-task learning to address these discrepancies, and predicts the drug response as the output. To the best of our knowledge, AITL is the first adversarial inductive transfer learning method to address both input and output discrepancies. Experimental results indicate that AITL outperforms state-of-the-art pharmacogenomics and transfer learning baselines and may guide precision oncology more accurately.

**Availability of codes and supplementary material:** https://github.com/hosseinshn/AITL

## 1 Introduction

The goal of pharmacogenomics (Evans and Relling, 1999) is to predict response to a drug given some single- or multi-omics data. Since clinical datasets in pharmacogenomics (patients) are small and hard to obtain, it is not feasible to train a computational model only on patients. As a result, many studies have focused on large pre-clinical pharmacogenomics datasets such as cancer cell lines as a proxy to patients (Barretina *et al.*, 2012; Iorio *et al.*, 2016). A majority of the current computational methods are trained on cell line datasets and then tested on other cell line or patient datasets (Sharifi-Noghabi *et al.*, 2019b; Sakellaropoulos *et al.*, 2019; Mourragui *et al.*, 2019; Rampášek *et al.*, 2019; Ding *et al.*, 2018; Geeleher *et al.*, 2017, 2014). However, cell lines and patients data, even with the same set of genes, do not have identical distributions due to the lack of an immune system and the tumor microenvironment in cell lines, which means a model cannot be trained on cell lines and then tested on patients (Mourragui *et al.*, 2019). Moreover, in cell lines, the response is often measured by the drug concentration that reduces viability by 50% (IC50), whereas in patients, it is often based on changes in the size of the tumor and measured by metrics such as response evaluation criteria in solid tumors (RECIST) (Schwartz *et al.*, 2016). This means that drug response prediction is a regression problem in cell lines but a classification problem in patients. As a result, discrepancies exist in both the input and output spaces in pharmacogenomics datasets. Therefore, a need exists for a novel method to address these discrepancies to utilize cell line and patient data together to build a more accurate model eventually for patients.

Transfer learning (Pan and Yang, 2009) attempts to solve this challenge by leveraging the knowledge in a *source* domain, a large data-rich dataset, to improve the generalization performance on a small *target* domain. Training a model on the source domain and testing it on the target domain violates the i.i.d assumption that the train and test data are from the same distribution. The discrepancy in the input space decreases the prediction accuracy on the test data, which leads to poor generalization (Zhang *et al.*, 2019). Transductive transfer learning (e.g. domain adaptation) and inductive transfer learning both use a labeled source domain to improve the generalization on a target domain. Transductive transfer learning assumes an unlabeled target domain, whereas inductive transfer learning assumes a labeled target domain where the label spaces of the source and target domain are different (Pan and Yang, 2009). Many methods have been proposed to minimize the discrepancy between the source and the target domains using different distribution metrics such as Maximum Mean Discrepancy (Gretton *et al.*, 2012) or using subspace-centric approaches such as Geodesic Flow Kernel (Gong *et al.*, 2012). In the context of drug response prediction, Mourragui *et al.* (Mourragui *et al.*, 2019) proposed PRECISE, a subspace-centric method, based on principal component analysis to minimize the discrepancy in the input space between cell lines and patients. Recently, adversarial domain adaptation has shown great performance in addressing the discrepancy in the input space for different applications, and its performance is comparable to the metric-based and subspace-centric methods in computer vision (Hosseini-Asl *et al.*, 2018; Pinheiro, 2018; Zou *et al.*, 2018; Tsai *et al.*, 2018; Long *et al.*, 2018; Chen *et al.*, 2017; Tzeng *et al.*, 2017; Ganin and Lempitsky, 2014). However, adversarial adaptation that addresses the discrepancies in both the input and output spaces has not yet been explored neither for pharmacogenomics nor for other applications.

In this paper, we propose Adversarial Inductive Transfer Learning (AITL), the first adversarial method of inductive transfer learning. Different from existing methods for inductive transfer learning as well as methods for adversarial transfer learning, AITL adapts not only the input space but also the output space. In pharmacogenomics, the source domain is the gene expression data obtained from the cell lines and the target domain is the gene expression data obtained from patients. Both domains have the same set of genes (i.e., raw feature representation). Discrepancies exist between the gene expression data in the input space, and the measure of the drug response in the output space. AITL learns features for the source and target samples and uses these features as input for a multi-task subnetwork to predict drug response for both the source and the target samples. The output space discrepancy is addressed by the multi-task subnetwork by assigning binary labels, called cross-domain labels, to the source samples which only have continuous labels. The multi-task subnetwork also alleviates the problem of small sample size in the target domain by joint training with the source domain. To address the discrepancy in the input space, AITL performs adversarial domain adaptation. The goal is that features learned for the source samples should be domain-invariant and similar enough to the features learned for the target samples to fool a global discriminator that receives samples from both domains. Moreover, with the cross-domain binary labels available for the source samples, AITL further regularizes the learned features by class-wise discriminators. A class-wise discriminator receives source and target samples from the same class label and should not be able to predict the domain accurately.

We evaluated the performance of AITL and state-of-the-art pharmacogenomics and transfer learning methods on pharmacogenomics datasets in terms of the Area Under the Receiver Operating Characteristic curve (AUROC) and the Area Under the Precision-Recall curve (AUPR). In our experiments, AITL achieved a substantial improvement compared to the baseline methods, demonstrating the potential of transfer learning for drug response prediction, a crucial task of precision oncology. Finally, we showed that the responses predicted by AITL for TCGA patients (without the drug response recorded) for breast, prostate, lung, kidney, and bladder cancers had statistically significant associations with the level of expression of some of the annotated target genes for the studied drugs. This shows that AITL captures biological aspects of the response.

## 2 Background and related work

### 2.1 Transfer Learning

Following the notation of (Pan and Yang, 2009), a domain like *DM* is defined by a raw input feature space^1^ **X**and a probability distribution *p*(*X*), where *X* = {*x*_1_, *x*_2_, &, *x_n_*} and *x_i_* is the *i*-th raw feature vector of *X*. A task *T* is associated with *DM* = {**X**, *p*(*X*)}, where *T* = {**Y**, *F* (.)} is defined by a label space **Y** and a predictive function *F* (.) which is learned from training data of the form (*X, Y*), where *X* ∈ **X** and *Y* ∈ **Y**. A labeled source domain is defined as 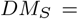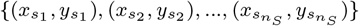 and a labeled target domain is defined as 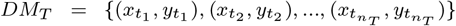, where *x_s_* ∈ *X_S_*, *x_t_* ∈ *X_T_*, *y_s_* ∈ *Y_S_*, and *y_t_* ∈ *Y_T_* . Since *n_T_* ≪ *n_S_*, it is challenging to train a model only on the target domain. Transfer learning addresses this challenge with the goal to improve the generalization on a target task *T_T_* using the knowledge in *DM_S_* and *DM_T_*, as well as their corresponding tasks *T_S_* and *T_T_* . Transfer learning can be categorized into three categories: 1) unsupervised transfer learning, 2) transductive transfer learning, and 3) inductive transfer learning. In unsupervised transfer learning, there is no label in the source and target domains. In transductive transfer learning, the source domain is labeled while the target domain is unlabeled. In this category, domains can be either the same or different (domain adaptation), but the source and target tasks are the same. In inductive transfer learning, the target domain is labeled and the source domain can be either labeled or unlabeled. In this category, the domains can be the same or different, but the tasks are always different (Pan and Yang, 2009).

### 2.2 Inductive transfer learning

There are three approaches to inductive transfer learning: 1) deep metric learning, 2) few-shot learning, and 3) weight transfer (Scott *et al.*, 2018). Deep metric learning methods are independent of the number of samples in each class of the target domain, denoted as *k*, meaning that they work for small and large *k* values. Few-shot learning methods focus on small *k* (*k* ≤ 20). Finally, weight transfer methods require a large k (*k* ≥ 100 or *k* ≥ 1000) (Scott *et al.*, 2018).

In drug response prediction, the target domain is small, which means a limited number of samples for each class is available; therefore, few-shot learning is more suitable for such a problem. Few-shot learning involves training a classifier to recognize new classes, provided only a small number of examples from each of these new classes in the training data (Snell *et al.*, 2017). Various methods have been proposed for few-shot learning (Chen *et al.*, 2019; Scott *et al.*, 2018; Snell *et al.*, 2017). For example, ProtoNet (Snell *et al.*, 2017) uses the source domain to learn how to extract features from the input and applies the feature extractor in the target domain. The mean feature of each class, obtained from the source domain, is used as the class prototype to assign labels to the target samples based on the Euclidean distance between a target sample’s feature and class prototypes.

### 2.3 Adversarial transfer learning

Recent advances in adversarial learning leverage deep neural networks to learn transferable representation that disentangles domain-invariant and class-invariant features from different domains and matches them properly (Peng *et al.*, 2019; Zhang *et al.*, 2019; Long *et al.*, 2018). In this section, we first introduce the Generative Adversarial Networks (GANs) (Goodfellow *et al.*, 2014), and then introduce some of the existing works on adversarial transfer learning.

#### 2.3.1 Generative Adversarial Networks

GANs (Goodfellow *et al.*, 2014) attempt to learn the distribution of the input *data* via a minimax framework where two networks are competing: a discriminator *D* and a generator *G*. The generator tries to create fake samples from a randomly sampled latent variable *z* that fool the discriminator, while the discriminator tries to catch these fake samples and discriminate them from the real ones. Therefore, the generator wants to minimize its error, while the discriminator wants to maximize its accuracy:

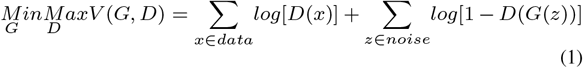

A majority of literature on adversarial transfer learning are for transductive transfer learning where the source domain is labeled while the target domain is unlabeled.

#### 2.3.2 Adversarial transductive transfer learning

Transductive transfer learning, often referred to as domain adaptation, is the most common scenario in transfer learning. Various methods have been proposed for adversarial transductive transfer learning in different applications such as image segmentation (Chen *et al.*, 2017), image classification (Tzeng *et al.*, 2017), speech recognition (Hosseini-Asl *et al.*, 2018), domain adaptation under label-shift (Azizzadenesheli *et al.*, 2019), and multiple domain aggregation (Schoenauer-Sebag *et al.*, 2019). The idea of these methods is that features extracted from source and target samples should be similar enough to fool a global discriminator (Tzeng *et al.*, 2017) and/or class-wise discriminators (Chen *et al.*, 2017).

## 3 Methods

### 3.1 Problem definition

Given a labeled source domain *DM_S_* with a learning task *T_S_* and a labeled target domain *DM_T_* with a learning task *T_T_*, where *T_T_* ≠ *T_S_*, and *p*(*X_T_*) ≠ *p*(*X_S_*), where *X_S_*, *X_T_* ∈ **X**, we assume that the source and the target domains are not the same due to different probability distributions. The goal of Adversarial Inductive Transfer Learning (AITL) is to utilize the source and target domains and their tasks in order to improve the learning of *F_T_* (.) on *DM_T_* .

In the area of pharmacogenomics, the source domain is the gene expression data obtained from the cell lines, and the source task is to predict the drug response in the form of IC50 values. The target domain consists of gene expression data obtained from patients, and the target task is to predict drug response in a different form – often change in the size of tumor after receiving the drug. In this setting, *p*(*X_T_*) ≠ *p*(*X_S_*) because cell lines are different from patients even with the same set of genes. Additionally, *Y_T_* ≠ *Y_S_* because for the target task *Y_T_* ∈ {0, 1}, drug response in patients is a binary outcome, but for the source task *Y_S_* ∈ *R*^+^, drug response in cell lines is a continuous outcome. As a result, AITL needs to address these discrepancies in both the input and output spaces.

### 3.2 AITL: Adversarial Inductive Transfer Learning

Our proposed AITL method takes input data from the source and target domains, and achieves the following three objectives: first, it makes predictions for the target domain using both of the input domains and their corresponding tasks, second, it addresses the discrepancy in the output space between the source and target tasks, and third, it addresses the discrepancy in the input space. AITL is a neural network consisting of four components:

- The feature extractor receives the input data from the source and target domains and extracts salient features, which are then sent to the multi-task subnetwork component.
- The multi-task subnetwork takes the extracted features of source and target samples and maps them to their corresponding labels and makes predictions for them. This component has a shared layer and two task-specific towers for regression (source task) and classification (target task). Therefore, by training the multi-task subnetwork on the source and target samples, it addresses the small sample size challenge in the target domain. In addition, it also addresses the discrepancy in the output space by assigning cross-domain labels (binary labels in this case) to the source samples (for which only continuous labels are available) using its classification tower.
- The global discriminator receives extracted features of source and target samples and predicts if an input sample is from the source or the target domain. To address the discrepancy in the input space, these features should be domain-invariant so that the global discriminator cannot predict their domain labels accurately. This goal is achieved by adversarial learning.
- The class-wise discriminators further reduce the discrepancy in the input space by adversarial learning at the level of the different classes, i.e., extracted features of source and target samples from the same class go to the discriminator for that class and this discriminator should not be able to predict if an input sample from a given class is from the source or the target domain.

The AITL cost function consists of a classification loss, a regression loss, and global and class-wise discriminator adversarial losses and is optimized end-to-end. An overview of the proposed method is presented in figure 1.

**Fig. 1.**
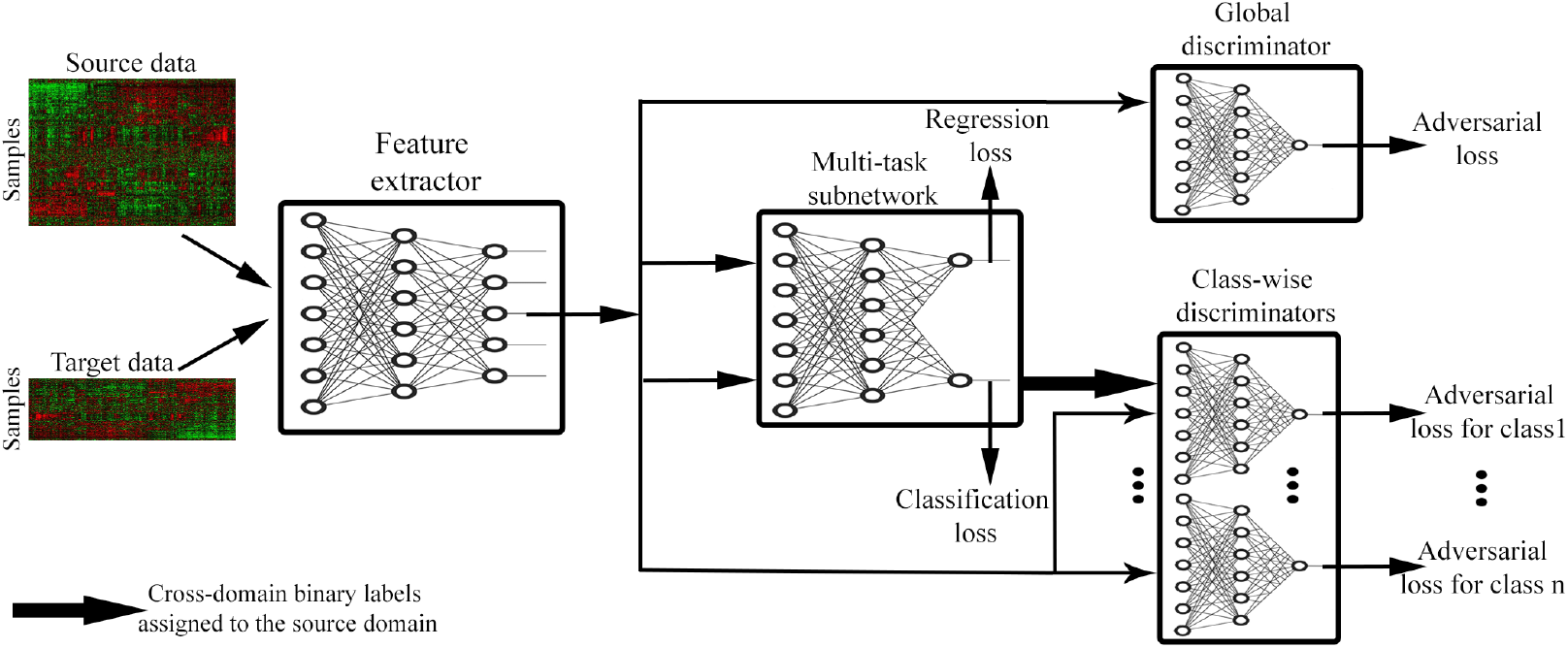
Schematic overview of AITL: First, the feature extractor receives source and target samples and maps them to a feature space in lower dimensions. Then, the multi-task subnetwork uses these features to make predictions for the source and target samples and also assigns cross-domain labels to the source samples. The multi-task subnetwork addresses the discrepancy in the output space. Finally, to address the input space discrepancy, global and class-wise discriminators receive the extracted features and regularize the feature extractor to learn domain-invariant features.

#### 3.2.1 Feature extractor

To learn salient features in lower dimensions for the input data, we design a feature extractor component. The feature extractor receives both the source and target samples as input and maps them to a feature space, denoted as *Z*. We denote the feature extractor as *f*(.):

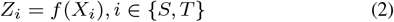

where *Z* denotes the extracted features for input *X* which is from either the source (*S*) or the target (*T*) domain. In pharmacogenomics, the feature extractor learns features for the cell line and patient data.

#### 3.2.2 Multi-task subnetwork

After extracting features of the input samples, we want to use these learned features to 1) make predictions for target samples, and 2) address the discrepancy between the source and the target domains in the output space. To achieve these goals, a multi-task subnetwork with a shared layer *g*(.) and two task-specific towers, denoted as *M_S_* (.) and *M_T_* (.), is designed, where *M_S_* is for regression (the source task) and *M_T_* is for classification (the target task):

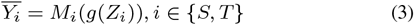

The performance of the multi-task subnetwork component is evaluated based on a binary-cross entropy loss for the classification task on the target samples and a mean squared loss for the regression task on the source samples:

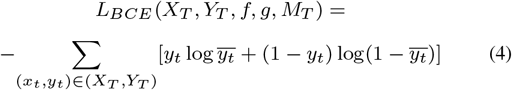

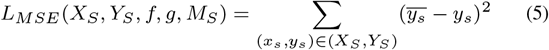

Where *Y_S_* and *Y_T_* are the true labels of the source and the target samples, respectively, and *L_BCE_* and *L_MSE_* are the corresponding losses for the target and the source domains, respectively. The multi-task subnetwork outputs 1) the predicted continuous labels for the source samples, 2) the predicted binary labels for the target samples, and 3) the assigned cross-domain binary labels for the source samples. The assigned cross-domain binary labels are obtained via the classification tower in the multi-task subnetwork which assigns binary labels (responder or non-responder) to the source samples because such labels do not exist for the source samples. Therefore, the multi-task subnetwork adapts the output space of the source and the target domains by assigning cross-domain labels to the source domain. In pharmacogenomics, the multi-task subnetwork predicts IC50 values for the cell lines and the binary response outcome for the patients. Moreover, it also assigns binary response labels to the cell lines which is similar to those of the patients.

#### 3.2.3 Global discriminator

The goal of this component is to address the discrepancy in the input space by adversarial learning of domain-invariant features. To achieve this goal, a discriminator receives source and target extracted features from the feature extractor and classifies them into their corresponding domain. The feature extractor should learn domain-invariant features to fool the global discriminator. In pharmacogenomics, the global discriminator should not be able to recognize if the extracted features of a sample are from a cell line or a patient. This discriminator is denoted as *D_G_*(.). The adversarial loss for *D_G_*(.) is as follows:

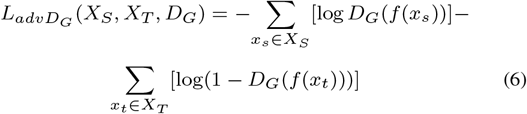

#### 3.2.4 Class-wise discriminators

With cross-domain binary labels available for the source domain, AITL further reduces the discrepancy between the input domains via class-wise discriminators. The goal is to learn domain-invariant features with respect to specific class labels such that they fool corresponding class-wise discriminators. Therefore, extracted features of the target samples in class *i*, and those of the source domain which the multi-task subnetwork assigned to class *i*, will go to the discriminator for class *i*. We denote such a class-wise discriminator as *DC_i_*. The adversarial loss for *DC_i_* is as follows:

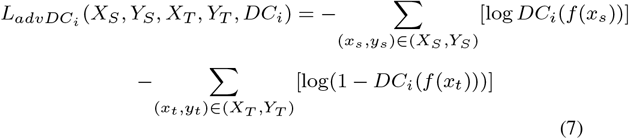

In pharmacogenomics, the class-wise discriminator for the responder samples should not be able to recognize if the extracted features of a responder sample are from a cell line or a patient (similarly for a non-responder sample).

#### 3.2.5 Cost function

To optimize the entire network in an end-to-end fashion, we design the cost function as follows:

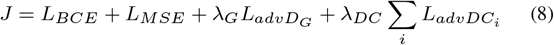

Where, *λ_G_* and *λ_DC_* are adversarial regularization coefficients for the global and class-wise discriminators, respectively.

#### 3.2.6 AITL architecture

The feature extractor is a one-layer fully-connected subnetwork with batch normalization using the ReLU activation function. The multi-task subnetwork has a shared fully-connected layer with batch normalization and the ReLU activation function. The regression tower has two layers (one hidden layer and one output layer) with the ReLU activation function in the first layer and the linear activation function in the second one. The classification tower has one fully-connected layer with the Sigmoid activation function that maps the features to the binary outputs directly. Finally, the global and class-wise discriminators are one-layer subnetworks with the Sigmoid activation function.

### 3.3 Drug response prediction for TCGA patients

To study AITL’s performance, similar to (Geeleher *et al.*, 2017; Sharifi-Noghabi *et al.*, 2019b), we employ the model trained on Docetaxel, Paclitaxel, or Bortezomib to predict the response for patients in several TCGA cohorts for which no drug response was recorded. For each drug, we extract the list of annotated target genes from the PharmacoDB resource (Smirnov *et al.*, 2017). We excluded Cisplatin because there was only one annotated target gene for it in PharmacoDB. To study associations between the level of expression of the extracted genes and the responses predicted by ATIL for each drug in each TCGA cohort, we fit multivariate linear regression models to the gene expression of those genes and the responses to that drug predicted by AITL. We obtain p-values for each gene and correct them for multiple hypothesis testing, using the Bonferroni correction (*α* = 0.05). The list of annotated target genes for each drug is available in the supplementary material (Section S1).

### 3.4 Datasets

In our experiments, we used the following datasets (See Table 1 for more detail):

- The Genomics of Drug Sensitivity in Cancer (GDSC) cell lines dataset, consisting of a thousand cell lines from different cancer types, screened with 265 targeted and chemotherapy drugs. (Iorio *et al.*, 2016)
- The Patient-Derived Xenograft (PDX) Encyclopedia dataset, consisting of more than 300 PDX samples for different cancer types, screened with 34 targeted and chemotherapy drugs. (Gao *et al.*, 2015)
- The Cancer Genome Atlas (TCGA) (Weinstein *et al.*, 2013) containing a total number of 117 patients with diverse cancer types, treated with Cisplatin, Docetaxel, or Paclitaxel (Ding *et al.*, 2016).
- Patient datasets from nine clinical trial cohorts containing a total number of 491 patients with diverse cancer types, treated with Bortezomib (Amin *et al.*, 2014; Mulligan *et al.*, 2007), Cisplatin (Silver *et al.*, 2010; Marchion *et al.*, 2011), Docetaxel (Hatzis *et al.*, 2011; Lehmann *et al.*, 2011; Chang *et al.*, 2005), or Paclitaxel (Hatzis *et al.*, 2011; Bauer *et al.*, 2010; Ahmed *et al.*, 2007).
- TCGA cohorts including, breast (BRCA), prostate (PRAD), lung (LUAD), kidney (KIRP), and bladder (BLCA) cancers that do not have the drug response outcome.

**Table 1.** Characteristics of the datasets

| Dataset | Resource | Drug | Type | Domain | Sample Size | Number of genes* |
| --- | --- | --- | --- | --- | --- | --- |
| GSE55145 (Amin *et al.*, 2014) | clinical trial | Bortezomib | targeted | target | 67 | 11609 |
| GSE9782-GPL96 (Mulligan *et al.*, 2007) | clinical trial | Bortezomib | targeted | target | 169 | 11609 |
| GDSC (Iorio *et al.*, 2016) | cell line | Bortezomib | targeted | source | 391 | 11609 |
| GSE18864 (Silver *et al.*, 2010) | clinical trial | Cisplatin | Chemotherapy | target | 24 | 11768 |
| GSE23554 (Marchion *et al.*, 2011) | clinical trial | Cisplatin | Chemotherapy | target | 28 | 11768 |
| TCGA (Ding *et al.*, 2016) | patient | Cisplatin | Chemotherapy | target | 66 | 11768 |
| GDSC (Iorio *et al.*, 2016) | cell line | Cisplatin | Chemotherapy | source | 829 | 11768 |
| GSE25065 (Hatzis *et al.*, 2011) | clinical trial | Docetaxel | Chemotherapy | target | 49 | 8119 |
| GSE28796 (Lehmann *et al.*, 2011) | clinical trial | Docetaxel | Chemotherapy | target | 12 | 8119 |
| GSE6434 (Chang *et al.*, 2005) | clinical trial | Docetaxel | Chemotherapy | target | 24 | 8119 |
| TCGA (Ding *et al.*, 2016) | patient | Docetaxel | Chemotherapy | target | 16 | 8119 |
| GDSC (Iorio *et al.*, 2016) | cell line | Docetaxel | Chemotherapy | source | 829 | 8119 |
| GSE15622 (Ahmed *et al.*, 2007) | clinical trial | Paclitaxel | Chemotherapy | target | 20 | 11731 |
| GSE22513 (Bauer *et al.*, 2010) | clinical trial | Paclitaxel | Chemotherapy | target | 14 | 11731 |
| GSE25065 (Hatzis *et al.*, 2011) | clinical trial | Paclitaxel | Chemotherapy | target | 84 | 11731 |
| PDX (Gao *et al.*, 2015) | animal (mouse) | Paclitaxel | Chemotherapy | target | 43 | 11731 |
| TCGA (Ding *et al.*, 2016) | patient | Paclitaxel | Chemotherapy | target | 35 | 11731 |
| GDSC (Iorio *et al.*, 2016) | cell line | Paclitaxel | Chemotherapy | source | 389 | 11731 |

\*Number of genes in common between the source and all of the target data for each drug

The GDSC dataset was used as the source domain, and all the other datasets were used as the target domain. For the GDSC dataset, raw gene expression data were downloaded from ArrayExpress (E-MTAB-3610) and response outcomes from https://www.cancerrxgene.org release 7.0. Gene expression data of TCGA patients were downloaded from the Firehose Broad GDAC (version published on 28.01.2016) and the response outcome was obtained from (Ding *et al.*, 2016). Patient datasets from clinical trials were obtained from the Gene Expression Omnibus (GEO), and the PDX dataset was obtained from the supplementary material of (Gao *et al.*, 2015). For each drug, we selected those patient datasets that applied a comparable measure of the drug response. For preprocessing, the same procedure was adopted as described in the supplementary material of (Sharifi-Noghabi *et al.*, 2019b) for the raw gene expression data (normalized and z-score transformed) and the drug response data. After the preprocessing, source and target domains had the same number of genes.

## 4 Results

### 4.1 Experimental design

We designed our experiments to answer the following four questions:

1. Does AITL outperform baselines that are trained only on cell lines and then evaluated on patients (without transfer learning)? To answer this question, we compared AITL against the method of (Geeleher *et al.*, 2014) and MOLI (Sharifi-Noghabi *et al.*, 2019b) which are state-of-the-art methods of drug response prediction that do not perform domain adaptation. The method of (Geeleher *et al.*, 2014) is non-deep learning method based on ridge regression and MOLI is a deep learning-based method. Both of them were originally proposed for pharmacogenomics.
2. Does AITL outperform baselines that adopt adversarial transductive transfer learning and non-deep learning adaptation (without adaptation of the output space)? To answer this question, we compared AITL against ADDA (Tzeng *et al.*, 2017) and the method of (Chen *et al.*, 2017), state-of-the-art methods of adversarial transductive transfer learning with global and class-wise discriminators, respectively. For the non-deep learning baseline, we compared AITL to PRECISE (Mourragui *et al.*, 2019), a non-deep learning domain adaptation method specifically designed for pharmacogenomics.
3. Does AITL outperform a baseline for inductive transfer learning? To answer this question, we compared AITL against ProtoNet (Snell *et al.*, 2017) which is a state-of-the-art inductive transfer learning method for small numbers of examples per class.
4. Finally, do the predicted responses by AITL for TCGA patients have associations with the targets of the studied drug?

Based on the availability of patient/PDX datasets for a drug, we experimented with four different drugs: Bortezomib, Cisplatin, Docetaxel, and Paclitaxel. It is important to note that these drugs have different mechanisms and are being prescribed for different cancers. For example, Docetaxel is a chemotherapy drug mostly known for treating breast cancer patients (Chang *et al.*, 2005), while Bortezomib is a targeted drug mostly used for multiple myeloma patients (Amin *et al.*, 2014). Therefore, the datasets we have selected cover different types of anti-cancer drugs.

In addition to the experimental comparison against published methods, we also performed an ablation study to investigate the impact of the different AITL components separately. AITL*−AD* denotes a version of AITL without the adversarial adaptation components, which means the network only contains the multi-task subnetwork. AITL*−D_G_* denotes a version of AITL without the global discriminator, which means the network only employs the multi-task subnetwork and class-wise discriminators. AITL*−DC* denotes a version of AITL without the class-wise discriminators, which means the network only contains the multi-task subnetwork and the global discriminator.

All of the baselines were trained on the same data, tested on patients/PDX for these drugs, and eventually compared to AITL in terms of prediction AUROC and AUPR. Since the majority of the studied baselines cannot use the continuous IC50 values in the source domain, binarized IC50 labels provided by (Iorio *et al.*, 2016) using the Waterfall approach (Barretina *et al.*, 2012) were used to train them. Finally, for the minimax optimization, a gradient reversal layer was employed by AITL and the adversarial baselines (Ganin *et al.*, 2016) which is a well-established approach in domain adaptation (Zhang *et al.*, 2019; Long *et al.*, 2018; You *et al.*, 2019). We performed 3-fold cross validation in the experiments to tune the hyper-parameters of AITL and the baselines based on the AUROC. Two folds of the source samples were used for training and the third fold for validation, similarly, two folds of the target samples were used for training and validation, and the third one for the test. The reported results refer to the average and standard deviation over the test folds. The hyper-parameters tuned for AITL were the number of nodes in the hidden layers, learning rates, mini-batch size, the dropout rate, number of epochs, and the regularization coefficients. We considered different ranges for each hyper-parameter and the final selected hyper-parameter settings for each drug and each method are provided in the supplementary material (Section S2). Finally, each network was re-trained on the selected settings using the train and validation data together for each drug. We used Adagrad for optimizing the parameters of AITL and the baselines (Duchi *et al.*, 2011) implemented in the PyTorch framework, except for the method of (Geeleher *et al.*, 2014) which was implemented in R. We used the author’s implementations for the method of (Geeleher *et al.*, 2014), MOLI, PRECISE, and ProtoNet. For ADDA, we used an existing implementation from https://github.com/jvanvugt/pytorch-domain-adaptation, and we implemented the method of (Chen *et al.*, 2017) from scratch.

### 4.2 Input and output space adaptation via AITL improves the drug response performance

Table 2 and Figure 2 report the performance of AITL and the baselines in terms of AUROC and AUPR, respectively. To answer the first experimental question, AITL was compared to the baselines which do not use any adaptation (neither the input nor the output space), i.e. the method of (Geeleher *et al.*, 2014) and MOLI (Sharifi-Noghabi *et al.*, 2019b), and AITL demonstrated a better performance in both AUROC and AUPR for all of the studied drugs. This indicates that addressing the discrepancies in the input and output spaces leads to better performance compared to training a model on the source domain and testing it on the target domain. To answer the second experimental question, AITL was compared to state-of-the-art methods of adversarial and non-deep learning transductive transfer learning, i.e. ADDA (Tzeng *et al.*, 2017), the method of (Chen *et al.*, 2017), and PRECISE (Mourragui *et al.*, 2019), which address the discrepancy only in the input space. AITL achieved significantly better performance in AUROC for all of the drugs and for three out of four drugs in AUPR (the results of (Chen *et al.*, 2017) for Cisplatin were very competitive with AITL). This indicates that addressing the discrepancies in the both spaces outperforms addressing only the input space discrepancy. Finally, to answer the last experimental question, AITL was compared to ProtoNet (Snell *et al.*, 2017) – a representative of inductive transfer learning with input space adaptation via few-shot learning. AITL outperformed this method in all of the metrics for all of the drugs.

**Table 2.** Performance of AITL and the baselines in terms of the prediction AUROC

| Method/Drug | Bortezomib | Cisplatin | Docetaxel | Paclitaxel |
| --- | --- | --- | --- | --- |
| (Geeleher et al., 2014) | 0.48 | 0.58 | 0.55 | 0.53 |
| MOLI (Sharifi-Noghabi et al., 2019b) | 0.57 | 0.54 | 0.54 | 0.53 |
| PRECISE (Mourragui et al., 2019) | 0.54 | 0.59 | 0.52 | 0.56 |
| (Chen et al., 2017) | 0.54±0.07 | 0.60±0.14 | 0.52±0.02 | 0.58±0.04 |
| ADDA (Tzeng et al., 2017) | 0.51±0.06 | 0.56±0.06 | 0.48±0.06 | did not converge |
| ProtoNet (Snell et al., 2017) | 0.49±0.01 | 0.40±0.003 | 0.40±0.01 | did not converge |
| AITL-AD<sup>1</sup> | 0.69±0.03 | 0.57±0.03 | 0.57±0.05 | 0.58±0.01 |
| AITL-D<sub>G</sub><sup>2</sup> | 0.69±0.04 | 0.62±0.1 | 0.48±0.03 | **0.62±0.02** |
| AITL-D<sub>C</sub><sup>3</sup> | 0.69±0.03 | 0.54±0.1 | 0.59±0.07 | 0.59±0.03 |
| AITL | **0.74±0.02** | **0.66±0.02** | **0.64±0.04** | 0.61±0.04 |

<sup>1</sup>AITL with only the multi-task subnetwork (no adversarial adaptation). <sup>2</sup>AITL with only class-wise discriminators and the multi-task subnetwork (no global discriminator). <sup>3</sup>AITL with only the global discriminator and the multi-task subnetwork (no class-wise discriminator)

**Fig. 2.**
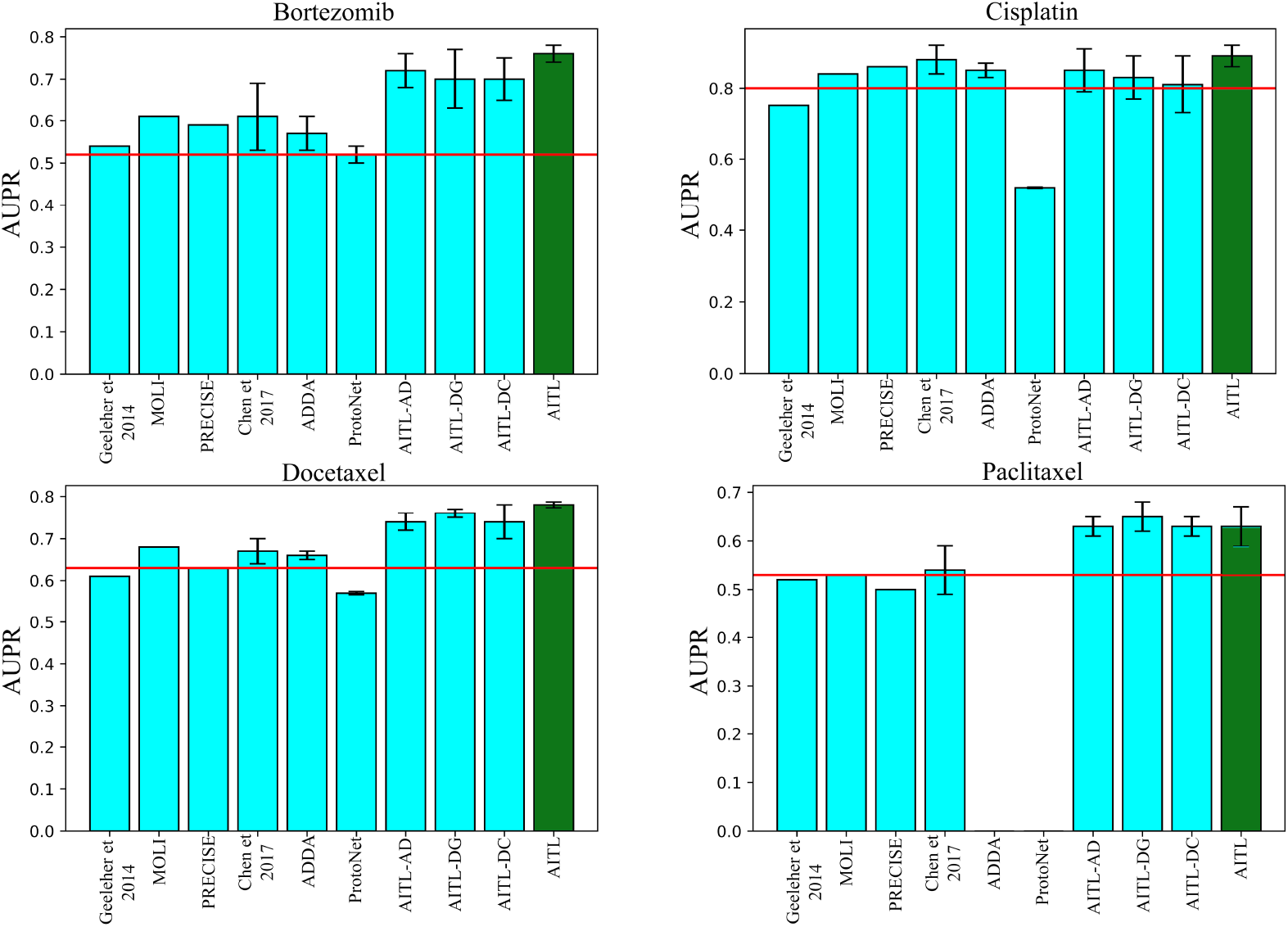
Performance of AITL and the baselines in terms of the prediction AUPR

We note that the methods of drug response prediction without adaptation, namely the method of (Geeleher *et al.*, 2014) and MOLI, outperformed the method of inductive transfer learning based on few-shot learning (ProtoNet). Moreover, these two methods also showed a very competitive performance compared to the methods of transductive transfer learning (ADDA, the method of (Chen *et al.*, 2017), and PRECISE). For Paclitaxel, ADDA did not converge in the first step (training a classifier on the source domain), which was also observed in another study (Sharifi-Noghabi *et al.*, 2019b). ProtoNet also did not converge for this drug.

We observed that AITL, using all of its components together, outperforms all the additional baselines omitting some of the components. This indicates the importance of both input and output space adaptation. The only exception was for the drug Paclitaxel, where AITL*−D_G_* outperforms AITL. We believe the reason for this is that this drug has the most heterogeneous target domain (see Table 1), and therefore the global discriminator component of AITL causes a minor decrease in the performance. All these results indicate that addressing the discrepancies in the input and output spaces between the source and target domains, via the AITL method, leads to a better prediction performance.

### 4.3 AITL predictions for TCGA patients have significant associations with target genes

To answer the last experimental question, we applied AITL models (trained on Docetaxel, Bortezomib, and Paclitaxel) to the gene expression data without known drug response from TCGA (breast, prostate, lung, kidney, and bladder cancers) and predicted the response for these patients separately. Based on the corrected p-values obtained from multiple linear regression, there are a number of statistically significant associations between the target genes of the studied drugs and the responses predicted by AITL. For example, in breast cancer, we observed statistically significant associations in MAP4 (*P* < 1 × 10^−^10) for Doxetaxel, BLC2 (*P* = 1.7 × 10^−^4) for Paclitaxel, and PSMA4 (*P* = 4.7 × 10^−^6) for Bortezomib. In prostate cancer, we observed statistically significant associations in MAP2 (*P*< 1 × 10^−^10) for Docetaxel, TUBB (*P* < 1 × 10^−^10) for Paclitaxel, and RELA (*P* = 2.2 × 10^−^4) for Bortezomib. For bladder cancer, NR1I2 (*P* = 0.04) for Docetaxel, MAP4 (*P* < 1 × 10^−^10) for Paclitaxel, and PSMA4 (*P* = 0.001) for Bortezomib were significant. In kidney cancer, BLC2 (*P* = 5.4 × 10^−^8) for Docetaxel, MAPT (*P* < 1 × 10^−^10) for Paclitaxel, and PSMD2 (*P* = 1 × 10^−^5) for Bortezomib were significant.

Finally, in lung cancer, MAP4 (*P* < 1 × 10^−^10) for Docetaxel, TUBB (*P* < 1 × 10^−^10) for Paclitaxel, and RELA (*P* < 1 × 10^−^10) for Bortezomib were significant. The complete list of statistically significant genes is presented in the supplementary material (Section S3).

The obtained results are in concordance with previous studies. For example, we observed that Microtubule-Associated Proteins (MAPs) were significant for Docetaxel and Paclitaxel in the studied cancers which aligns with previous research on this family of proteins (Yang *et al.*, 2017; Smoter *et al.*, 2011; Bhat and Setaluri, 2007). For Bortezomib, we observed significant associations for different proteasome subunits such as subunit alpha (PSMA) and beta (PSMB). These subunits have been shown to be key players across different cancers (Rouette *et al.*, 2016; Tsvetkov *et al.*, 2017; Li *et al.*, 2017). We also observed significant associations for RELA (also known as Transcription Factor p65) in all of the studied cancers which aligns with its oncogenic role across different cancers (Collignon *et al.*, 2018), and moreover, with its reported associations with Bortezomib in breast cancer (Hideshima *et al.*, 2014), prostate cancer (Manna *et al.*, 2013), and lung cancer (Zhao *et al.*, 2015). All these results suggest that AITL predictions capture biological aspects of the drug response.

### 4.4 Discussion

To our surprise, ProtoNet, and ADDA could not outperform the method of (Geeleher *et al.*, 2014), MOLI, and PRECISE. For ProtoNet, this may be due to the depth of the backbone network. A recent study has shown that a deeper backbone improves ProtoNet performance significantly in image classification Chen *et al.* (2019). However, in pharmacogenomics, employing a deep backbone is not realistic because of the much smaller sample size compared to an image classification application. Another limitation for ProtoNet is the imbalanced number of training examples in different classes in pharmacogenomics datasets. Specifically, the number of examples per class in the training episodes is limited to the number of samples of the minority class as ProtoNet requires the same number of examples from each class. For ADDA, this lower performance may be due to the lack of end-to-end training of the classifier along with the global discriminator of this method. The reason is that end-to-end training of the classifier along with the discriminators improved the performance of the second adversarial baseline (Chen *et al.*, 2017) in AUROC and AUPR compared to ADDA. Moreover, the method of (Chen *et al.*, 2017) also showed a relatively better performance in AUPR compared to the method of (Geeleher *et al.*, 2014) and MOLI.

In pharmacogenomics, patient datasets with drug response are small or not publicly available due to privacy and/or data sharing issues. We believe including more patient samples and more drugs will increase generalization capability. In addition, recent pharmacogenomics studies have shown that using multi-omics data works better than using only gene expression (Sharifi-Noghabi *et al.*, 2019b). In this work, we did not consider genomic data other than gene expression data due to the lack of patient samples with multi-omics and drug response data publicly available; however, in principle, AITL also can work with such data. Last but not least, we used pharmacogenomics as our motivating application for this new problem of transfer learning, but we believe that AITL can also be employed in other applications. For example, in slow progressing cancers such as prostate cancer, large patient datasets with gene expression and short-term clinical data (source domain) are available, however, patient datasets with long-term clinical data (target domain) are small. AITL may be beneficial to learn a model to predict these long-term clinical labels using the source domain and its short-term clinical labels (Sharifi-Noghabi *et al.*, 2019a).

For future research directions, we believe that the TCGA dataset consisting of gene expression data of more than 12,000 patients (without drug response outcome) can be incorporated in an unsupervised transfer learning setting to learn better features that are domain-invariant between cell lines and cancer patients. The advantage of this approach is that we can keep the valuable patient datasets with drug response as an independent test set and not use it for training/validation. Another possible future direction is to incorporate domain-expert knowledge into the structure of the model. A recent study has shown that such a structure improves the drug response prediction performance on cell line datasets and, more importantly, provides an explainable model as well (Snow *et al.*, 2019).

## Acknowledgements

We would like to thank Hossein Asghari, Baraa Orabi, and Yen-Yi Lin (the Vancouver Prostate Centre) and Soufiane Mourragui (the Netherlands Cancer Institute) for their support. We also would like to thank the Vancouver Prostate Centre for providing the computational resources for this research.

## Authors’ contributions

Study concept and design: H.SN., C.C.C., M.E. Deep learning design, implementations, and analysis: H.SN. and S.P. Data preprocessing, analysis, and interpretation: O.Z. Analysis and interpretation of results: H.SN., S.P., O.Z. Drafting of the manuscript: All authors read and approved the final manuscript. Supervision: C.C.C., M.E.

## 5 Conclusion

In this paper, we introduced a new problem in transfer learning motivated by applications in pharmacogenomics. Unlike domain adaptation that only requires adaptation in the input space, this new problem requires adaptation in both the input and output spaces.

To address this problem, we proposed AITL, an Adversarial Inductive Transfer Learning method which, to the best of our knowledge, is the first method that addresses the discrepancies in both the input and output spaces. AITL uses a feature extractor to learn features for target and source samples. Then, to address the discrepancy in the output space, AITL utilizes these features as input of a multi-task subnetwork that makes predictions for the target samples and assign cross-domain labels to the source samples. Finally, to address the input space discrepancy, AITL employs global and class-wise discriminators for learning domain-invariant features. In pharmacogenomics, AITL adapts the gene expression data obtained from cell lines and patients in the input space, and also adapts different measures of the drug response between cell lines and patients in the output space. In addition, AITL can also be employed in other applications such as predicting long-term clinical labels for slow progressing cancers.

We evaluated AITL on four different drugs and compared it against state-of-the-art baselines in terms of AUROC and AUPR. The empirical results indicated that AITL achieved a significantly better performance compared to the baselines showing the benefits of addressing the discrepancies in both the input and output spaces. Finally, we analyzed AITL’s predictions for the studied drugs on breast, prostate, lung, kidney, and bladder cancer patients in TCGA. We showed that AITL’s predictions have statistically significant associations with the level of expression of some of the annotated target genes for the studied drugs. We conclude that AITL may be beneficial in pharmacogenomics, a crucial task in precision oncology.

## Footnotes

1 This is different from learned features by the network

